# Tissue-specific inhibition of protein sumoylation uncovers diverse SUMO functions during *C. elegans* vulval development

**DOI:** 10.1101/2021.12.06.471357

**Authors:** Aleksandra Fergin, Gabriel Boesch, Nadja R. Greter, Simon Berger, Alex Hajnal

**Author notes:** Molecular Life Science PhD Program, University and ETH Zürich, CH-8057 Zürich, Switzerland.

## Abstract

The sumoylation (SUMO) pathway is involved in a variety of processes during *C. elegans* development, such as gonadal and vulval fate specification, cell cycle progression and maintenance of chromosome structure. The ubiquitous expression of the sumoylation machinery and its involvement in many essential processes has made it difficult to dissect the tissue-specific roles of protein sumoylation and identify the specific target proteins. To overcome these challenges, we have established tools to block protein sumoylation and degrade sumoylated target proteins in a tissue-specific and temporally controlled manner. We employed the auxin-inducible protein degradation system (AID) to down-regulate AID-tagged SUMO E3 ligase GEI-17 or the SUMO ortholog SMO-1, either in the vulval precursor cells (VPCs) or in the gonadal anchor cell (AC). Tissue-specific inhibition of GEI-17 and SMO-1 revealed diverse roles of the SUMO pathway during vulval development, such as AC positioning, basement membrane (BM) breaching, vulval cell fate specification and epithelial morphogenesis. Inhibition of sumoylation in the VPCs resulted in an abnormal shape of the vulval toroids and ectopic cell fusions. Sumoylation of the ETS transcription factor LIN-1 at K169 mediates a subset of these SUMO functions, especially the proper contraction of the ventral vulA toroids. Thus, the SUMO pathway plays diverse roles throughout vulval development.

## Introduction

Sumoylation is an essential post-translational protein modification found in eukaryotes (Matunis, Wu, and Blobel 1998; Mahajan et al. 1997). A major player in this pathway is the so-called Small Ubiquitin-like Modifier (SUMO), which shares large structural and functional similarities with Ubiquitin. However, unlike ubiquitination sumoylation promotes or inhibits protein interactions and changes protein conformation or localization, allowing transient binding to target proteins. Since its discovery in the late 1990s, SUMO has been shown to be involved in a wide range of essential biological processes (Johnson 2004; Hay 2005; Wilkinson and Henley 2010; Flotho and Melchior 2013). Though, studying its diverse functions and identifying specific targets has remained challenging due to the essential roles of the SUMO pathway for animal viability and development, the low concentration of sumoylated target proteins, the constant activity of deSUMOylating enzymes and the often subtle effects caused by the modification itself (Johnson 2004; Flotho and Melchior 2013). Developing tools, which allow spatial and temporal inhibition of protein sumoylation, is therefore crucial to gain a better understanding of this protein modification.

Protein sumoylation in *C. elegans* occurs in essentially the same fashion as in higher organisms. However, contrary to mammals and other vertebrates, only one SUMO orthologue, called SMO-1, exists in *C. elegans*. This renders the worm an ideal model to study this post-translational protein modification system. Activated SMO-1 is transferred to the E2 enzyme UBC-9 by the E1 enzyme formed by UBA-2 and AOS-1, and attached to the substrate by a SUMO E3 ligase, such as the PIAS domain protein GEI-17 (Holway, Hung, and Michael 2005; Ferreira et al. 2013; Pelisch et al. 2014). Deconjugation of SUMO from its targets is regulated by one of four SUMO proteases, ULP-1, ULP-2, ULP-4 and ULP-5 (Pelisch et al. 2014).

Sumoylation is essential for *C. elegans* viability and involved in a wide range of biological processes. Particularly, vulval development has previously been shown to depend on sumoylation, as *smo-1(lf)* mutants exhibit multivulva (Muv) as well as protruding vulva (Pvl) phenotypes (Broday 2004). We therefore chose this well-established model to further dissect the different roles of the SUMO pathway during organogenesis (Schindler and Sherwood 2013; Schmid and Hajnal 2015). The vulva is formed by three out of six equivalent vulval precursor cells (the VPCs P3.p trough P8.p), which adopt one of three possible cell fates. The 1° fate is induced in P6.p by an epidermal growth factor (EGF) signal, termed LIN-3, which is secreted by the gonadal anchor cell (AC) 11/30/21 2:17:00 PM. A lateral signal from P6.p then activates the LIN-12 Notch signaling pathway in the neighboring VPCs P5.p and P7.p to induce the alternate, secondary (2°) fate (Greenwald, Sternberg, and Horvitz 1983; Greenwald 1985; Sundaram 2004; Greenwald 2005). After vulval fate specification, the 1° VPC undergoes three rounds of cell divisions producing 8 daughter cells, while the 2° fated VPCs each generate 7 daughter cells in an asymmetric lineage, together forming the vulva consisting of 22 cells. The remaining distal VPCs (P3.p, P4.p and P8.p) adopt the uninduced 3° fate, which is to divide once and fuse with the surrounding epidermis hyp7. While the VPCs proliferate, the AC breaches two basement membranes (BMs) separating the uterus from the epidermis and invades the underlying vulval epithelium (Sherwood and Sternberg 2003). During the subsequent phase of vulval morphogenesis, the vulval cells invaginate to generate a lumen, extend circumferential protrusions and fuse with their contralateral partner cells to form a tubular organ consisting of a stack of seven epithelial rings called toroids (Schindler and Sherwood 2013; Schmid and Hajnal 2015).

Here, we have employed a tissue-specific version of the auxin-inducible protein degradation system (AID) to inhibit the SUMO pathway either in the AC or the vulval cells (Zhang et al. 2015). This approach allowed us to determine, in which tissues protein sumoylation is necessary for normal vulval development, as well as to characterize the diverse phenotypes caused by selectively blocking the SUMO pathway. Moreover, we hypothesized that, by inserting a degron tag into SUMO itself, we could induce degradation of sumoylated proteins in a tissue-specific manner. To test if this approach allows the in vivo identification of SUMO targets, we chose LIN-1 as it was previously shown to be sumoylated at K10 and K169 by in vitro experiments (Leight 2005; Leight et al. 2015). The ETS family transcription factor LIN-1 is essential for different aspects of vulval development. During fate specification, LIN-1 inhibits VPC differentiation by recruiting transcriptional repressors in a sumoylation-dependent and independent manner to repress 1° fate-specific target genes (Miley et al. 2004; Leight 2005), and during morphogenesis, LIN-1 promotes the contraction of ventral toroids (Farooqui et al. 2012).

We mutated the two known sumoylation sites in LIN-1 (K10 and K169), measured LIN-1 expression levels after VPC-specific degradation of AID-tagged SMO-1 and observed the vulval phenotypes caused by mutation of the SUMO sites. This approach suggested that sumoylation of LIN-1 in the VPCs at K169 is required for the proper contraction of the ventral vulA toroid ring during morphogenesis. Additional phenotypes that were only observed after degradation of GEI-17 or SMO-1 suggest that LIN-1 is one of several relevant SUMO targets during vulval development.

This approach may be used to investigate the in vivo significance of potential SUMO targets identified through in vitro experiments or by proteomic methods.

## Results

### Tissue-specific, auxin-inducible degradation of SUMO pathway components

To dissect the interactions of different SUMO pathway components with their substrates and identify specific targets during vulval development, we adapted the tissue-specific auxin-inducible degradation system (Zhang et al. 2015). Here, we generated tissue-specific degradation drivers expressing the TIR-1 ubiquitin ligase in four different cell types; *hlh-2p>tir-1* in the AC and VU cells before and after AC specification (Sallee and Greenwald 2015), *cdh-3p>tir-1* in the AC post specification (Sherwood and Sternberg 2003), *egl-17p>tir-1* in the 1° VPC and its descendants (Burdine, Branda, and Stern 1998) and *bar-1p>tir-1* in all VPCs and their descendants (Eisenmann et al. 1998). We also used an existing driver, in which TIR-1 is ubiquitously expressed under the *eft-3p>tir-1* promoter in somatic tissues (Zhang et al. 2015).

Specificity of the degradation drivers was assessed with three assays. First, we used an SL2 trans-splicing acceptor to express the mCherry fluorophore under the same promoter/enhancer as TIR-1. In this way tissue-specificity could be monitored by observing mCherry expression (**Fig. 1A**). Second, we assessed the loss of target protein expression by degrading a GFP- and AID-double tagged variant of GEI-17 (GFP::AID::GEI-17) (Pelisch et al. 2014). A strong decrease in GFP::AID::GEI-17 expression upon auxin treatment was only observed in tissues expressing TIR-1 (**Fig. 1B**). Lastly, we confirmed protein degradation by Western blot analysis in animals expressing the pan-somatic TIR-1 driver, treated with auxin for 1h or 24h (**Fig. S1A, B**). We detected only residual amounts of GEI-17 protein post degradation with comparable levels for both treatment periods. The residual levels most likely stem form GEI-17 expressed in the germline, where the TIR-1 driver was not expressed. Interestingly, degradation of GEI-17 did not affect expression levels of SMO-1 (**Fig. S1A, C**), suggesting that depletion of GEI-17 does not significantly alter the levels of free SMO-1.

**Fig 1.**
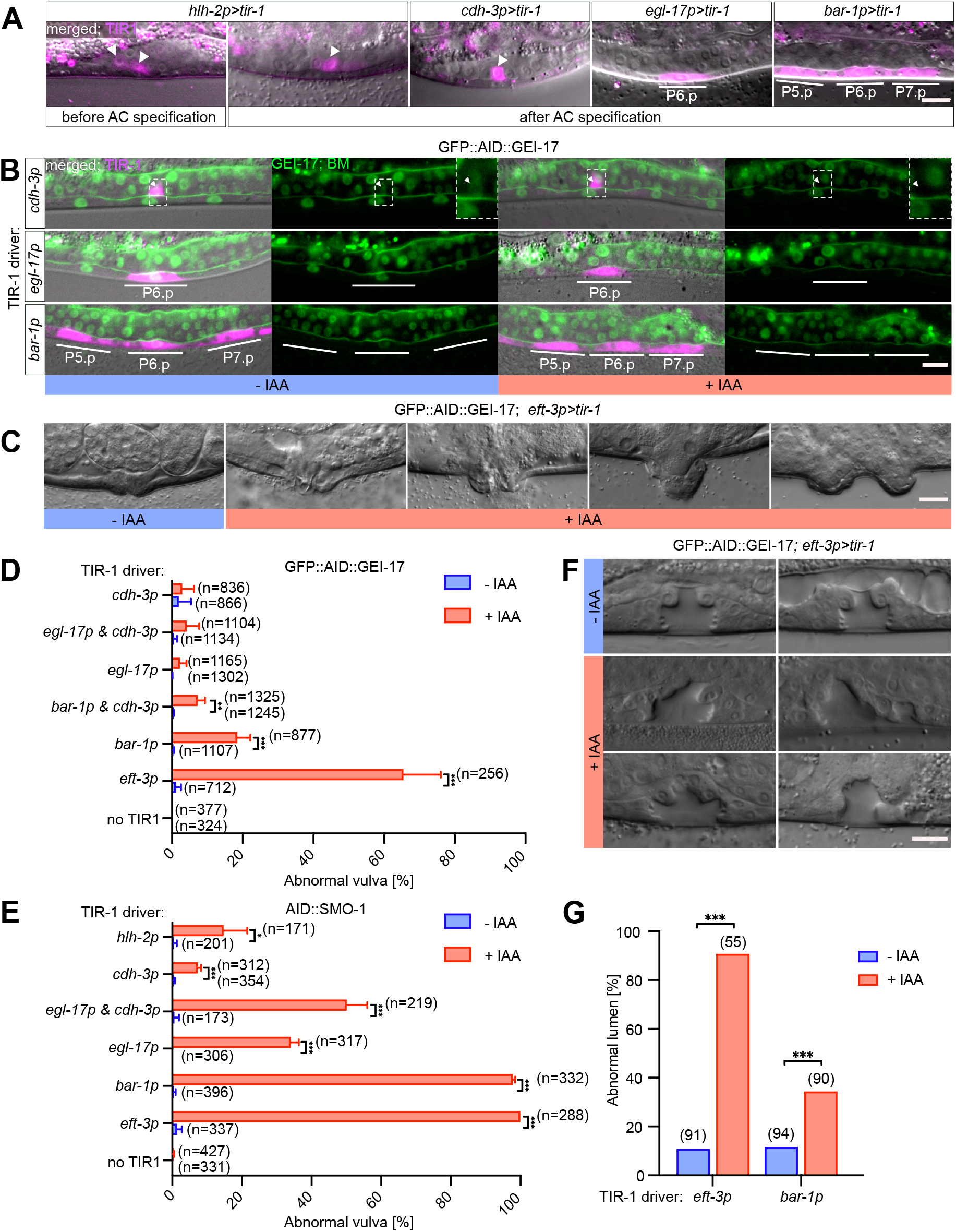
Degradation of SUMO pathway components leads to abnormal vulval development. **(A)** Tissue-specific expression of the TIR-1::SL2::mCherry degradation driver using the indicated TIR-1 drivers. mCherry expression from the bi-cistronic mRNA in magenta overlaid with the corresponding DIC images is shown for each transgene. White arrowheads indicate the AC and bars outline the location of the VPCs. **(B)** Tissue-specific degradation of GFP::AID::GEI-17 after auxin treatment using the indicated TIR-1 drivers. Left panels show the TIR-1::SL2::mCherry expression in magenta overlaid with the GFP::AID::GEI-17 and LAM-1::GFP BM markers in green and the corresponding DIC images. Right panels show only the GFP::AID::GEI-17 signal along with the LAM-1::GFP marker in green. White lines outline the location of the VPCs. The insets in the top row show the region around the AC magnified around 3x, indicated by white arrowhead. **(C)** DIC images illustrating the vulval morphology defects in adults after global degradation of GFP::AID::GEI-17. **(D)** Penetrance of the vulval morphology defects after auxin-induced degradation of GFP::AID::GEI-17 and (**E**) AID::SMO-1 using the indicated TIR-1 drivers. The mean values ± s.d. obtained from three biological replicates are shown. **(F**) DIC images of L4 larvae showing an abnormally shaped vulval lumen resulting after global GFP::AID::GEI-17 degradation. (**G**) Penetrance of the vulval morphogenesis defects shown in **(F)** using the global *eft-3p* and VPC-specific *bar-1p*>*tir-1* drivers. Treatment conditions are indicated as +IAA (blue) for animals treated with 1 mM auxin, and –IAA (red) for control animals. All GFP::AID::GEI-17 animals were treated at 25 °C from the L1 stage onward. All AID::SMO-1 were treated at 20 °C from the L2 to L4 stage. In **(D)** and **(E)** the numbers of animals scored are indicated in brackets. Statistical significance was determined by two-tailed unpaired t-tests **(D, E)** or with Mann-Whitney tests **(G)**. Asterisks indicate the p-values as * p≤0.05; ** p≤ 0.01; *** p≤ 0.001. The scale bars are 10 μm.

### Inhibiting the SUMO pathway in the VPCs or AC causes abnormal vulval development

Following the initial validation of our approach, we assessed how degradation of SUMO pathway components affects vulval development. For this purpose, we crossed the different TIR-1 degradation drivers with the GFP::AID::GEI-17 strain (*gei-17(fgp1)*, Pelisch et al. 2014) and an N-terminally tagged AID::SMO-1 allele (*smo-1(zh140)*, this study). Note that homozygous *smo-1(zh140)* animals showed a wild-type vulval morphology, but were sterile as adults. Degradation of either protein using the pan-somatic TIR-1 driver resulted in characteristic vulval morphogenesis defects (shown for GFP::AID::GEI-17 in **Fig. 1C**), similar to chromosomal mutations in SUMO pathway genes (Broday 2004). Most adult animals showed a protruding vulva (Pvl) or abnormal eversion (Evl) phenotype (Seydoux, Salvage, and Greenwald 1993) of varying severity and penetrance (**Fig. 1C-E**).

Degrading AID::SMO-1 in all somatic cells using the *eft-3p>tir-1* driver resulted in nearly completely penetrant vulval defects, while ubiquitous GEI-17 degradation caused abnormal vulval development in only 65.5% (SD ± 10.5) of the animals (**Fig. 1D, E**). Pan-somatic degradation caused generally more penetrant defects than tissue-specific degradation, except for depletion of AID::SMO-1 with the VPC-specific (*bar-1p>tir-1*) driver, which resulted in almost fully penetrant vulval defects (97.9% SD ± 0.6) (**Fig. 1E**). The combination of the AC (*cdh-3p>tir-1*) and 1° VPC (*egl-17p>tir-1*) -specific drivers resulted in an additive effect (50.2% SD ± 5.8 combined versus 7.4% SD ± 0. 9 and 34.1% SD ± 2.2 separate, respectively), suggesting that the observed morphogenesis defects are a combination of separate functions played by SMO-1 in those two tissues. By contrast, degradation of GFP::AID::GEI-17 with the VPC-specific driver resulted in less penetrant defects (18.5% SD ± 3.6), and degradation using the AC and 1° VPC drivers alone or in combination did not result in significant defects (**Fig. 1D**).

We further investigated the defects caused by GEI-17 depletion on vulval lumen formation in L4 larvae, after the toroids have been formed and the connection between vulva and uterus been established (**Fig. 1F, G**). After somatic GEI-17 degradation with *eft-3p>tir-1*, 90 % animals exhibited a misshaped vulval lumen, possibly due to defects in toroid fusion, cell migration defects or a failure to connect the vulva to the uterus. VPC-specific degradation using *bar-1p>tir-1*, on the other hand, had a less pronounced effects with only 34 % of the animals showing abnormal vulval morphogenesis (**Fig. 1F, G**), suggesting that the SUMO pathway is not only necessary in the VPCs but also in other tissues.

The overall lower penetrance of vulval defects observed after degrading GFP::AID::GEI-17 compared to AID::SMO-1 may be explained by the facts that SMO-1 is the only known SUMO orthologue in *C. elegans*, whereas GEI-17 is not the only E3 ligase, and that not all sumoylation reactions require an E3 ligase (Rai et al. 2011).

### The SUMO pathway acts during all stages of vulval development

To determine the developmental stage, at which sumoylation is required for proper vulval development, we degraded SMO-1 by exposing animals to auxin at varying developmental time points between the L1/2 and L3/4 molts (**Fig. S2A**), or by withdrawing auxin at different time points (**Fig. S2B**) and assessing the penetrance of the observed vulval defects.

In case of the *eft-3p>tir-1* and *bar-1p>tir-1* drivers, both an early auxin treatment during L1 until the L2 molt or a late treatment beginning in L3 caused highly penetrant vulval defects. Even though there may be a slight delay until the auxin-induced effect fades after removing the animals from auxin-containing medium (Zhang et al. 2015), these data point to a continuous action of the SUMO pathway throughout vulval development, from VPC fate specification until lumen morphogenesis.

### The SUMO pathway regulates VPC fate specification

VPC fate specification occurs between the late L2 and early L3 stages and requires the combined action of the Delta/Notch and EGRF/RAS/MAPK signaling pathways (Sundaram 2004). We first examined how inhibition of the SUMO pathway through VPC-specific degradation of GEI-17 affects 1° VPC fate specification. The *egl-17* gene, which encodes an FGF-like growth factor, can serve as a specific marker for the 1° VPC fate induced in reponse to EGRF/RAS/MAPK signaling (Burdine, Branda, and Stern 1998). We thus analyzed the expression of a transcriptional *egl-17>yfp* reporter after auxin-induced degradation of AID::GEI-17 with the *bar-1p>tir-1* driver. *egl-17>yfp* expression was stongly reduced in the 1° VPC P6.p and in its descendants at the two-(Pn.px) and four-cell (Pn.pxx) stages (**Fig. 2A, B**). The SUMO pathway therefore positively regulates 1° VPC fate specification.

**Fig 2.**
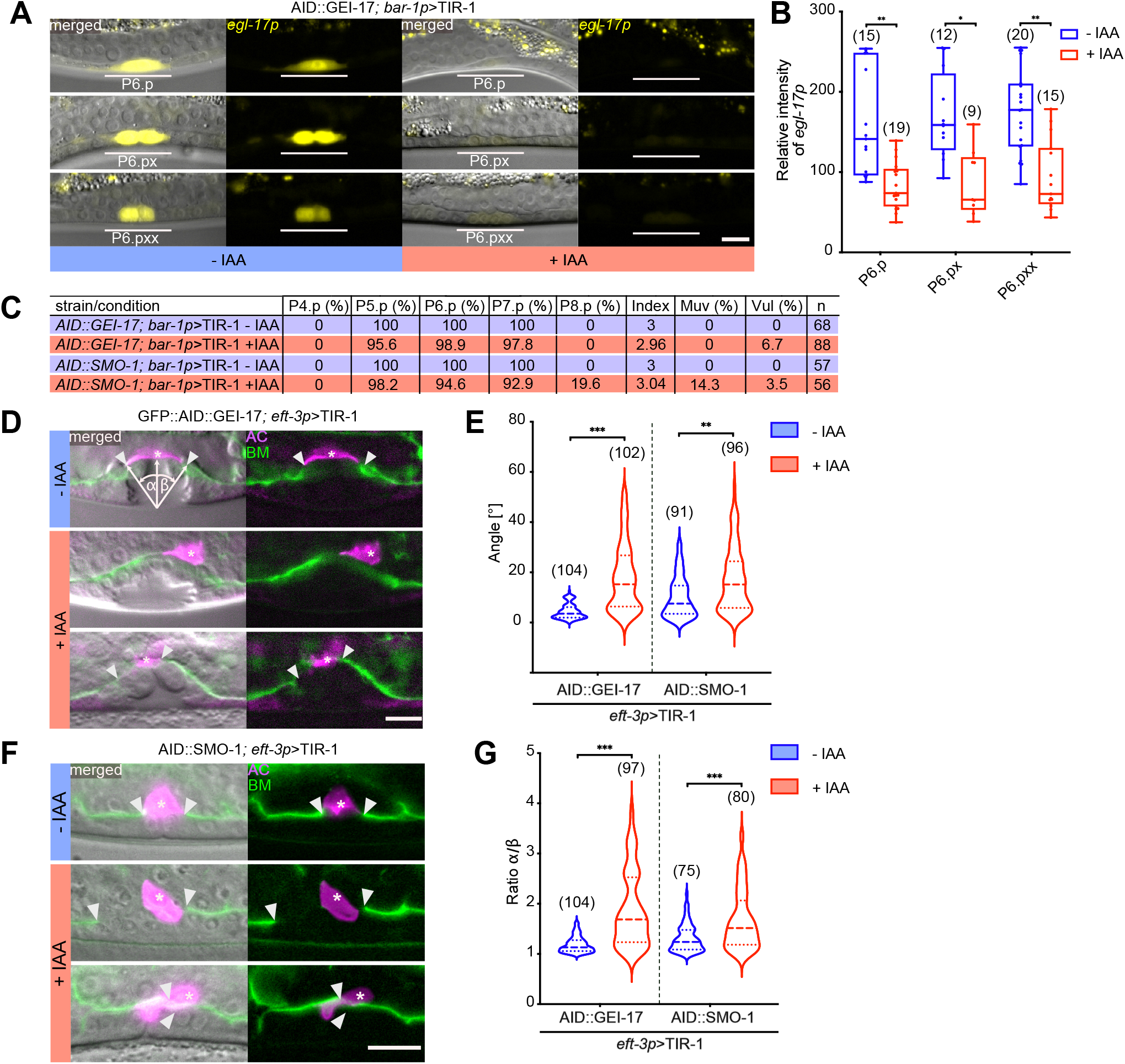
The SUMO pathway regulates vulval development. **(A)** *egl-17p>yfp* expression in P6.p and its descendants P6.px and P6.pxx after VPC-specific degradation of GEI-17. **(B)** Quantification of *egl-17p>yfp* expression levels after AID::GEI-17 depletion. Box plots show the median values with the 25^th^ and 75^th^ percentiles and whiskers indicate the maximum and minimum values. **(C)** VPC induction upon degradation of AID::GEI-17 and AID::SMO-1. For each strain and condition, the percent of induced VPCs, the average number of induced VPCs per animal (index), percent of multivulva (Muv, index>3) and vulvaless (Vul, index <3) animals, and the number of animals scored (n) are shown. **(D)** AC displacement, AC fusion defects and asymmetric BM breaching after global AID::GEI-17 and **(F)** AID::SMO-1 degradation. The BMs are labelled with LAM-1::GFP in green and the AC with *cdh-3p*>mCherry::moeABD in magenta. White arrowheads indicate the borders of the BM breaches and asterisks the AC. The left panels show the fluorescent signals merged with the corresponding DIC images. The angles α and β used to quantify AC alignment and symmetry of the BM breaching are illustrated in the top left panel. **(E)** Quantification of the AC displacement and **(G)** BM breaching asymmetry after degradation of GEI-17 and SMO-1 using the global *eft-3p>tir-1* driver. See also **suppl. Fig. S3** for the results obtained with tissue-specific *tir-1* drivers. Dashed lines in the violin plots **(E, G)** show the median values and the dotted lines the 25^th^ and 75^th^ percentiles. In all experiments, untreated controls are labelled with –IAA (blue) and animals treated with 1 mM auxin +IAA (red). In each graph, the numbers of animals scored are indicated in brackets. Statistical significance was determined with a Kolmogorov-Smirnov test **(B, E, G)**. p-values are indicated as * p≤0.05; ** p≤ 0.01; *** p≤ 0.001. The scale bars are 10 μm.

Poulin et al. (2004) and Broday et al. (2005) previously reported that a global loss of protein sumoylation in *smo-1(lf)* mutants or by *smo-1* RNAi caused the ectopic induction of additional VPCs besides the three proximal VPCs (P5.p to P7.p), leading to a multivulva (Muv) phenotype (Broday 2004; Poulin et al. 2005). To further quantify VPC fate specification after degradation of the SUMO pathway components, we counted the numbers of induced VPCs per animal after VPC-specific degradation of AID::GEI-17 (*zh142*, a *gei-17* allele containing an AID but no GFP tag) or AID::SMO-1 using the *bar-1p>tir-1* driver. Vulval induction after auxin-induced depletion of GEI-17 was slightly decreased (6.7% Vul, 2.96 VPCs/animal induced), consistent with the reduced levels of *egl-17>yfp* (**Fig. 2A-C**). Degradation of AID::SMO-1, on the other hand, resulted in a mixed phenotype with 14.3% of the animals showing ectopic induction and 3.5% an underinduced phenotype, but overall only a very slightly hyper-induced phenotype (3.04 VPCs/animal induced). Interestingly, we only observed ectopic induction of the posterior VPC P8.p, but never of the two anterior VPCs P3.p, P4.p (**Fig. 2C**).

Together, these data indicated that protein sumoylation in the VPCs both promotes the induction of the three proximal VPCs and inhibits the differentiation of the posterior VPC P8.p.

### Sumoylation is required for proper AC positioning and symmetrical BM breaching

Next, we analyzed the effects of SUMO pathway by examining AC positioning as well as BM breaching. The AC in animals globally depleted of AID::GFP::GEI-17 often failed to invade at the vulval midline and sometimes did not breach the BMs or breached them in an asymmetric fashion (**Fig. 2D**). In addition, the AC did not fuse in 69% of the animals to form the uterine seam cell syncytium (utse), which connects the vulva to the uterus, (**Fig. 2D** and **Fig. S3D**). In many cases, the AC was not properly positioned at the vulval midline (quantified in **Fig. 2E** as the angle of deflection from the midline), which may have led to the asymmetric or absent BM breaching (quantified in **Fig. 2G**). Mispositioning of the AC and BM breaching defects were only observed after somatic, but not after VPC-or AC-specific AID::GFP::GEI-17 degradation (**Fig. S2A, B**), suggesting that signals from additional tissues besides the VPCs control AC positioning (Ihara et al. 2011). Somatic degradation of SMO-1 also caused AC mispositioning and asymmetric BM breaching (**Fig. 2E-G**). As for GEI-17, neither VPC-nor AC-specific degradation of SMO-1 resulted in AC positioning or BM breaching defects (**Fig. S2A, B**)

In summary, our data indicate that the SUMO pathway is necessary for proper AC positioning and symmetrical BM breaching during invasion. This function appears to depend on a non-autonomous function of the SUMO pathway in tissues other than the AC and VPCs.

### The SUMO pathway is required for proper toroid morphogenesis

Since virtually all SMO-1 and most GEI-17-depleted animals showed abnormal vulval development as adults (**Fig. 1D, E**), while the VPC fate specification defects were comparably rare (**Fig. 2C**), we speculated that the inhibition of protein sumoylation perturbs vulval development predominantly during the later stage of morphogenesis. To characterize vulval morphogenesis in more detail, we therefore examined the structure of the vulval toroids. To monitor toroid formation, we used either the AJM-1::GFP or the HMR-1::GFP reporter, which both label the adherens junctions between the vulval cells (Köppen et al. 2001; Marston et al. 2016). Degradation of SMO-1 or GEI-17 in the VPCs using the *bar-1p>tir-1* driver lead to a number of different defects in toroid morphology. Specifically, we observed an abnormal shape of the ventral vulA toroids (**Fig. 3A**) and ectopic fusion between the vulC and vulD or the vulA and vulB1 toroids (arrows in **Fig. 3A**), similar to the defects observed in *smo-1* null mutants (Broday 2004). During normal vulval morphogenesis, the ventral toroids formed by the 2° VPCs contract in order to extend the apical lumen dorsally (Farooqui et al. 2012). To quantify ventral toroid contraction, we measured the ratio of the vulA to vulB1 diameters (**Fig. 3B**). The elevated vulA/vulB1 ratio indicated that the vulA toroids did not fully contract after inhibition of the SUMO pathway in the VPCs.

**Fig. 3.**
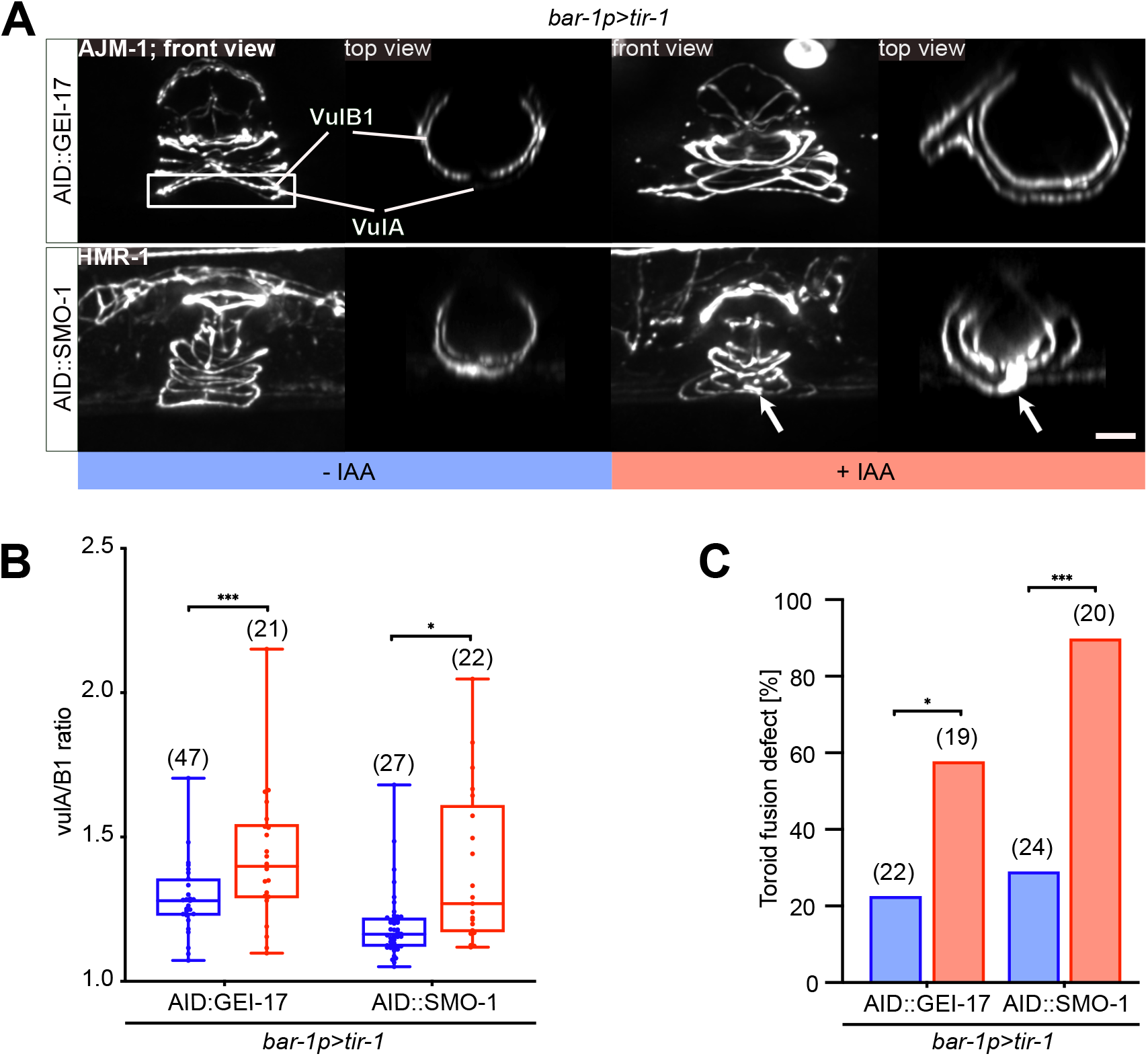
Inhibition of the SUMO pathway in the VPCs causes toroid morphogenesis defects. **(A)** Toroid morphogenesis defects in L4 hermaphrodites. 3D reconstructions of the adherens junctions labelled with AJM-1::GFP (for AID::GEI-17) or HMR-1::GFP (for AID::SMO-1) after VPC-specific degradation. Left panels show lateral views of z-projections. vulA and vulB1 toroids are outlined by the white rectangle in the top left panel and shown in top (xz) views in the right panels. White arrows point to abnormal fusion between the vulA and vulB1 toroids after AID::SMO-1 degradation. **(B)** Quantification of vulA contraction, calculated as the ratio of the vulA and vulB1 toroid diameter after VPC-specific AID::GEI-17 or AID::SMO-1 degradation. The box plots show the median values with the 25^th^ and 75^th^ percentiles and the whiskers indicate the maximum and minimum values. **(C)** Penetrance of toroid fusion defects after VPC-specific AID::GEI-17 or AID::SMO-1 degradation. In all experiments, untreated controls are labelled with –IAA (blue) and animals treated with 1 mM auxin +IAA (red). In each graph, the numbers of animals scored are indicated by the numbers in brackets. In **(B)** unpaired two-tailed t-tests and in **(C)** Mann-Whitney tests were used to determine statistical significance. p-values are indicated as * p≤0.05; ** p≤ 0.01; *** p≤ 0.001. The scale bar is 10 μm.

Taken together, these data indicated that the SUMO pathway acts in the VPCs during vulval toroid morphogenesis.

### Sumoylation stabilizes the LIN-1 protein in the 1° VPCs

The ETS family transcription factor LIN-1 is necessary to inhibit VPC fate specification during vulval induction and for the contraction of the ventral toroids during vulval morphogenesis (Miley et al. 2004; Leight 2005; Farooqui et al. 2012). To assess the role of LIN-1 sumoylation in vivo, we generated point mutations in the endogenous *lin-1* locus by replacing the two lysine residues K10 and K169 in the SUMO consensus motifs with alanine residues (Leight 2005; Leight et al. 2015). To monitor effects on LIN-1 expression levels, the two SUMO site mutations were introduced into the *lin-1(st12212)* background, in which a *gfp* tag had been inserted at the *lin-1* C-terminus. The wild-type *lin-1(st12212)* as well as the *lin-1(zh159)* K10A, K169A double mutant reporter were then crossed with the AID::SMO-1 allele and the VPC-specific *bar-1p>tir-1* driver. Wild-type LIN-1::GFP protein expression levels decreased in the 1° VPC descendants, once AID::SMO-1 was degraded through addition of auxin (**Fig. 4A**,, **Fig. S3A, B**). In untreated animals, LIN-1::GFP expression levels in the 2° VPC descendants were lower than in the 1° cells, and LIN-1::GFP expression in the 2° cells decreased only slightly after degradation of AID::SMO-1. Interestingly, the expression levels of the double mutant LIN-1(K10A, K169A)::GFP protein were already lower in the 1° cells of untreated animals, suggesting that sumoylation stabilizes the LIN-1 protein. Auxin-induced degradation of AID::SMO-1 did not cause a further decrease in LIN-1(K10A, K169A)::GFP levels in the 1° cells, indicating that the SUMO site mutations render LIN-1::GFP resistant to AID::SMO-1-mediated degradation. In the 2° VPC descen ants, however, a slight decrease in LIN-1(K10A, K169A)::GFP levels was observed after AID::SMO-1 degradation, suggesting that the sumoylation pathway may indirectly regulate LIN-1 levels in the 2° cells (**Fig. 4A, B, Fig. S3 A, B**).

**Fig. 4.**
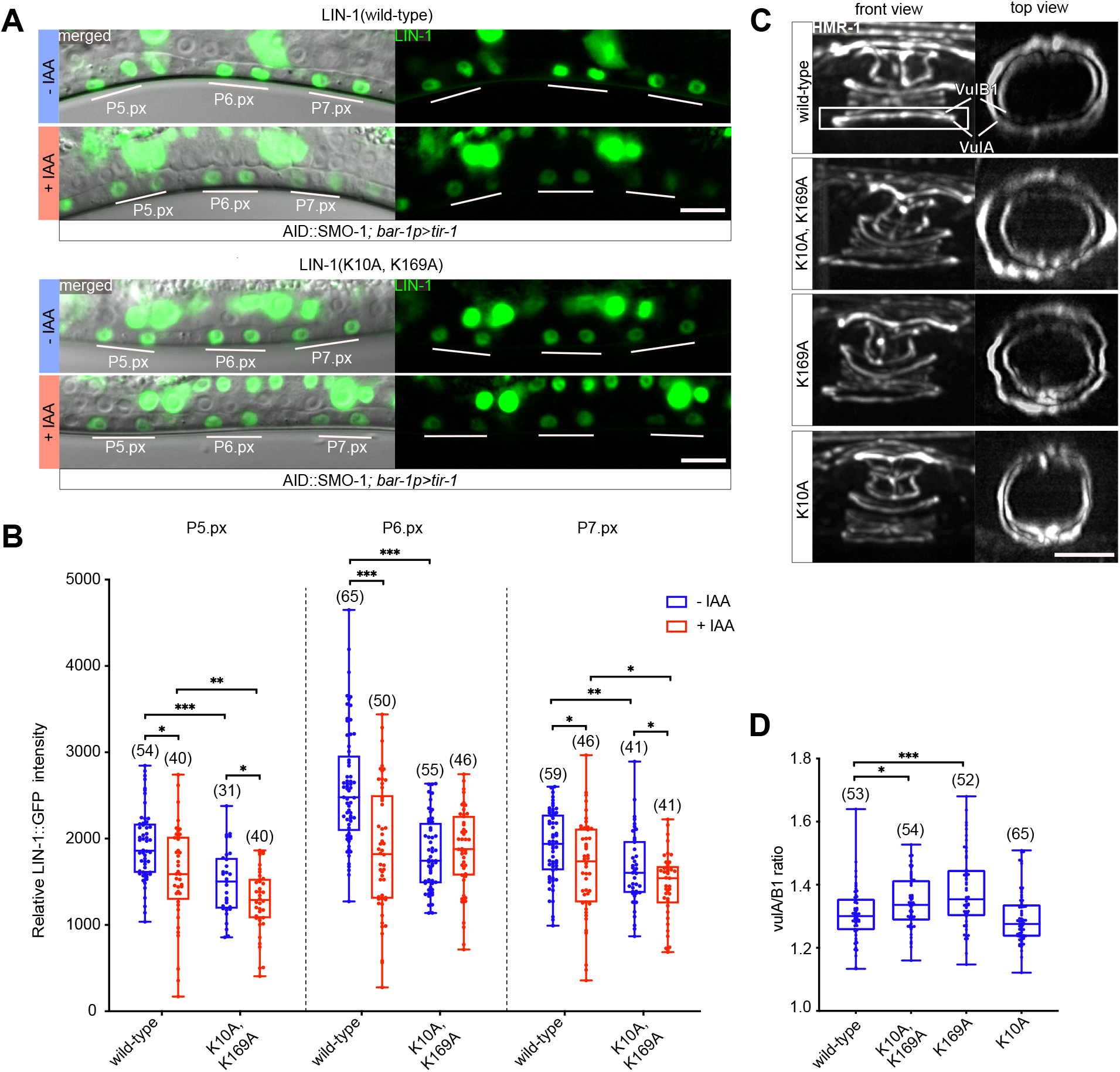
LIN-1 sumoylation is required for ventral toroid contraction. **(A)** Wild-type and K10A, K169A mutant LIN-1::GFP expression in L3 larvae at the Pn.px stage after VPC-specific degradation of AID::SMO-1 from the L2 stage onward. The 1° and 2° VPC descendants are underlined in white. The left panels show the corresponding DIC images overlaid with the LIN-1::GFP signal in green. **(B)** Quantification of LIN-1::GFP expression levels in 1° and 2° VPC descendants at the Pn.px stage in LIN-1::GFP wild-type and K10A, K169A double mutants under the indicated conditions. See **suppl. Fig. S4** for the corresponding measurements at the Pn.pxx stage. **(C)** Toroid morphogenesis defects in LIN-1 K10A and K169A single and double mutants at the L4 stage. Left panels show lateral views of z-projections. vulA and vulB1 toroids are outlined by the white rectangle in the top left panel and shown in top (xz) views in the right panels. **(D)** Quantification of vulA contraction, calculated as the ratio of the vulA and vulB1 toroid diameter. The box plots show the median values with the 25^th^ and 75^th^ percentiles and the whiskers indicate the maximum and minimum values. Where indicated, untreated controls are labelled with –IAA (blue) and animals treated with 1 mM auxin with +IAA (red). In each graph, the numbers of animals scored are indicated by the numbers in brackets. Statistical significance in **(B)** and (**D**) was calculated with unpaired two-tailed t-tests. p-values are indicated as * p≤0.05; ** p≤ 0.01; *** p≤ 0.001. The scale bars are 10 μm.

The reduced expression of wild-type, but not K10A, K169A double mutant LIN-1::GFP after AID::SMO-1 degradation suggested that a substantial fraction of endogenous LIN-1 is sumoylated in the 1° VPC descendants. These data indicated that proteasomal degradation of SUMO via AID can lead to the simultaneous degradation of SUMO-modified target proteins such as LIN-1. Furthermore, since LIN-1(K10A, K169A)::GFP levels were already reduced in the absence of auxin when compared to wild-type LIN-1::GFP, sumoylation may stabilize the LIN-1 protein. Possibly, only sumoylated LIN-1 can interact with certain binding partners to form a stable complex (Leight et al. 2015). Though, we cannot exclude the possibility that the K10A, K169A mutations may also affect other post-translational modifications of LIN-1, such as acetylation, methylation or ubiquitination, which could also affect LIN-1 stability and activity.

### The LIN-1 K169 SUMO site is necessary for ventral toroid contraction

To investigate the relevance of the two SUMO sites in LIN-1, we introduced the HMR-1::GFP adherens junction marker into *lin-1* single and double SUMO site mutants and investigated toroid formation (*lin-1(zh157)* and *lin-1(zh158)* refer to the K10A and K169A single SUMO site mutants, respectively.). In *lin-1(zh159)* K10A, K169A double and *lin-1(zh158)* K169A single mutants, we observed similar toroid contraction defects as seen after AID::SMO-1 or AID::GEI-17 degradation. Specifically, the ratio of the vulA to vulB1 diameter was increased in *zh159* double and *zh158* single mutants (**Fig. 4C, D**). By contrast, we did not detect any toroid morphogenesis defects in *lin-1(zh157)* K10A single mutants.

Thus, only the K169 SUMO site appears to be relevant for a specific aspect of LIN-1 function during vulval toroid morphogenesis. While sumoylation of LIN-1 at K169 may be required for the proper contraction of the ventral vulA toroids, none of the other defects observed after inhibition of the SUMO pathway were detected in the LIN-1 SUMO site mutants. The other functions of the SUMO pathway thus appear to be independent of LIN-1 sumoylation.

## Discussion

### A tissue-specific degradation toolkit to study the SUMO pathway and its targets

Posttranslational protein modification via the SUMO pathway is essential for many biological processes (Deyrieux and Wilson 2017). Though, the pleiotropic effects and transient and reversible nature of protein sumoylation has rendered this pathway difficult to study. The identification of SUMO targets is usually performed by proteomic approaches (Matunis and Rodriguez 2016; Hendriks and Vertegaal 2016) or through in vitro experiments, but the possibilities to validate candidate SUMO substrates in vivo have so far been limited.

Here, we applied the auxin-inducible protein degradation system AID (Zhang et al. 2015) to inactivate the SUMO pathway in a tissue-specific and temporally controlled manner. This approach may allow the verification of relevant SUMO targets in tissues of interest by following their expression levels after AID::SUMO degradation and observing the resulting phenotypes.

Using the *C. elegans* vulva as a model for organogenesis, we dissected the role of the SUMO pathway by inducing degradation of endogenously AID-tagged alleles of the SUMO homolog SMO-1 and the SUMO E3 ligase GEI-17 in the different cell types contributing to the vulva. For this purpose, we generated four tissue-specific TIR-1 driver lines to induce AID in the tissues of interest. Tissue-specificity was validated through fluorescent co-expression markers, degradation of a fluorescently tagged SUMO E3 ligase as well as protein quantification, confirming the effectiveness of the toolkit. The inducible nature of the AID system allowed us to assess the spatial and temporal requirements for protein sumoylation at different stages of vulval development. The stronger penetrance of defects observed after degrading SMO-1 in the VPCs compared to the AC suggested that sumoylation is predominantly required in the VPCs. This observation is consistent with previously reported roles of SMO-1 during vulval development (Miley et al. 2004; Leight et al. 2015; Ward et al. 2013). Temporally controlled depletion of SMO-1 indicated that protein sumoylation is continuously required throughout vulval development, controlling a variety of processes like VPC fate specification, AC positioning, BM breaching and vulval toroid morphogenesis. These findings expand the range of previously reported SUMO phenotypes and provide new insights in the role of sumoylation during vulval development.

Overall, global degradation of either SMO-1 or GEI-17 resulted in stronger and more penetrant phenotypes than AC- and VPC-specific degradation. This suggested that the SUMO pathway acts in additional tissues besides the AC and VPCs to control vulval development. For example, neurons in the ventral nerve cord are known to secrete AC guidance cues (Ziel et al. 2009), while adjacent muscles can affect VPC fate specification (Moghal 2003). Degradation of SMO-1 in general caused more severe phenotypes than GEI-17 degradation. This could be due to the fact that SMO-1 is the only *C. elegans* SUMO homolog, while GEI-17 is one of several known E3 ligases. GEI-17 may for example be replaced by MMS-21 (Rai et al. 2011), and sumoylation may even occur without an E3 ligase (Gareau and Lima 2010). Moreover, proteasomal degradation of AID-tagged SMO-1 appears to lead to the simultaneous degradation of sumoylated target proteins, as shown here for the case of LIN-1. This may be another factor explaining the stronger phenotypes observed after SMO-1 degradation compared to GEI-17 depletion.

The identification of SUMO targets is usually performed by proteomic approaches (Matunis and Rodriguez 2016; Hendriks and Vertegaal 2016) or through in vitro experiments, but the possibilities to validate candidate SUMO substrates in vivo have so far been limited. The tissue-specific AID approach presented here may allow the verification of relevant SUMO targets in specific cell types by following their expression levels after AID::SMO-1 degradation and observing the resulting phenotypes.

### The SUMO pathway is required for proper BM breaching by the AC

After specification of the VPC fates and before the onset of vulval morphogenesis, the AC breaches two BMs separating the uterus from the vulval cells and invades at the vulval midline in between the 1° vulF cells. Degradation of SMO-1 or GEI-17 resulted in characteristic AC invasion defects. Occasionally, the AC completely failed to breach the underlying BMs, but a more frequent defect was the displacement of the AC from the vulval midline, leading to asymmetric BM breaching. Global but neither VPC-nor AC-specific degradation of GEI-17 or SMO-1 resulted in a displacement of the AC and asymmetrical BM breaching. However, we were not able to pin-point the tissue, in which sumoylation affects AC positioning. After AC invasion, the VPCs continue to proliferate and invaginate, thereby enlarging the breach in the BM. The BMs then slide over the dividing vuF and vuE cells and are stabilized over then un-divided vulD cells, where the INA-1/PAT-3 integrins and the VAB-19 adhesion protein are expressed (Ding 2003; Ihara et al. 2011). BM sliding also depends on ventral uterine cells adjacent to the AC. LIN-12 Notch signaling in the uterine π cells upregulates *ctg-1* expression, which allows BM sliding by downregulating the dystroglycan BM-adhesion receptor (McClatchey et al. 2016). As reported by Broday et al. (2004) and consistent with our observations, sumoylation is required for the formation of a uterine lumen. The abnormal connection between the vulva and uterus may in part be caused by a loss of sumoylation of LIN-11 at K17 and K18 and a disruption of its function in π cells. AC positioning, on the other hand, depends on guidance signals from both the VPCs and the ventral nerve cord that polarize the AC along the dorso-ventral axis (Ziel et al. 2009; Naegeli et al. 2017). We thus speculate that the mispositioning of the AC and asymmetrical BM breaching are caused by a cumulative effect of multiple defects in different tissues.

### The SUMO pathway in the VPCs controls vulval toroid morphogenesis

The most penetrant class of phenotypes caused by disruption of the SUMO pathway affects vulval toroid morphogenesis. All toroid morphogenesis defects could be observed with the *bar-1p>tir-1* driver, indicating that these phenotypes are likely due to a cell-autonomous function of the SUMO pathway in the vulval cells. Inhibiting the SUMO pathway altered *egl-17* gene expression in the dividing 1° VPCs, already before the morphogenesis phase, indicating an involvement of SUMO pathway during VPC induction (Leight 2005, 1; Ward et al. 2013; Leight et al. 2015). Moreover, we did observe rare defects in proximal VPC induction and an ectopic induction of the posterior VPC P8.p after inhibition of the SUMO pathway, which also points to a role in VPC fate specification. Even though we could not directly correlate VPC induction with AC positioning, it is possible that the ectopic VPC induction is at least in part due to the AC mispositioning. However, the observed vulval morphogenesis defects were almost fully penetrant, indicating that the SUMO pathway is most relevant during morphogenesis, after the VPC fates have been specified. Protein sumoylation is required for different aspects of vulval morphogenesis, such as the formation of the correct connections and fusion between the contralateral pairs of vulval cells and for the contraction of the ventral vulA toroids.

### LIN-1 sumoylation site at K169 promotes the contraction of the ventral vulval toroids

The ETS family transcription factor LIN-1 is a well-characterized SUMO target, originally identified in genetic screens for mutants with abnormal vulval development (Miley et al. 2004; Leight 2005; Leight et al. 2015). While a complete loss of *lin-1* function causes a completely penetrant Muv phenotype due to loss of its repressor function, *lin-1(lf)* mutations also cause reduced *egl-17* reporter expression in the 1° VPC lineage (Tiensuu et al. 2005). Moreover, LIN-1 promotes ventral toroid contraction by inducing expression of the RHO kinase LET-502 in the 2° toroids (Farooqui et al. 2012). Together, these findings suggested that an inhibition of LIN-1 sumoylation could be responsible for a subset of the similar defects caused by inhibition of the SUMO pathway. Consistent with this hypothesis, expression levels of wild-type LIN-1 were reduced after degradation of AID::SMO-1, suggesting that LIN-1 is indeed sumoylated in the vulval cells. Moreover, deletion of the K169 sumoylation site in LIN-1 caused similar defects in vulA toroid contraction as VPC-specific inhibition of the SUMO pathway. We thus propose that sumoylation of LIN-1 at K169 is necessary for this specific activity during vulval toroid formation. Vulval fate specification, on the other hand, was not affected by deletion of either of the two SUMO sites in LIN-1.

In conclusion, our findings point to complex interactions between the SUMO pathway and various targets, depending on cellular context and developmental stage.

## Material and Methods

### *C. elegans* handling and maintenance

*Caenorhabditis elegans* strains were grown on standard NGM (Nematode Growth Medium) plates seeded with OP50 *E. coli* bacteria and incubated at 15 °C, 20 °C or 25 °C as indicated (Brenner 1974). The derivate of Bristol strain N2 was used as a wild-type reference. A list of strains used in this study is provided in **suppl. Table 1**.

### Design of the tissue-specific degradation toolkit

All TIR-1 degradation drivers were designed with an analogous design in the pCFJ151 backbone and integrated by MosSCI in selected genetic locations. To track the tissue-specificity of each construct, we used an SL2 trans-splicing domain followed by an mCherry reporter (fragment derived from pSA120 (Armenti et al. 2014)) to express the fluorophore under the same promoter as TIR-1. In all constructs, we used the *unc-54* 3’ UTR. TIR-1 was amplified from pLZ31 (Zhang et al. 2015). The following promoters/enhancers were used: the *egl-17* promoter was amplified as a 2042 bp fragment from a derivate of pPD107.94/mk84-148 (Kirouac and Sternberg 2003), the *cdh-3* promoter was amplified as a 1897 bp fragment from a derivate of pPD104.97/mk62-63 (Kirouac and Sternberg 2003; Ziel et al. 2009), the *bar-1* promoter as 3216 bp fragment (Nusser-Stein et al. 2012) and the *hlh-2prox* promoter as a 576 bp fragment driving the expression in two alpha and two beta cells (3VU and 1AC) (Sallee and Greenwald 2015). The promoters/enhancers are indicted in text as *egl-17p, cdh-3p, bar-1p* and *hlh-2p*. All constructs were cloned by Gibson assembly (Gibson et al. 2009; Gibson 2011). The following plasmid constructs were microinjected at the indicated final concentrations into young adult EG6699, EG8078 or EG8080 hermaphrodites: transgene in pCFJ150: 50 ng/μl, transformation markers pGH8 (*rap-3p>mCherry*): 10 ng/μl, pCFJ104 (*myop-3>mCherry*): 5 ng/μl, pCFJ90 (*myo-2p>mCherry*): 2.5 ng/μl and pJL43.1 expressing Mos1 transposase: 50 ng/μl (Frøkjær-Jensen et al. 2008). The transformants were screened for crawling animals, which lacked the co-injected transformation markers and genotyped for homozygous insertion by PCR. The list of plasmids generated and primers used for amplification of selected fragments and genotyping can be found in **suppl. Tables 2 & 3** in the **Suplementary material**.

### CRISPR/Cas9 genome editing

For CRISPR/Cas9 editing, the protocol by Dickinson et al. (2015) was followed. Plasmids containing the repair template and single guide RNAs were used at a concentration of 10 ng/μl and 50 ng/μl, respectively. We used the same transformation markers at the same concentrations as for MosSCI insertions. To generate the *smo-1(zh140)* allele, an oligonucleotide corresponding to a target sequence near the *smo-1* translational start site (sgRNA: GCC GAT GAT GCA GCT CAA GC) was cloned into the plasmid pMW46 (derivate of pDD162 from Addgene). The 5’homology arm was amplified from genomic DNA with OAF239 and OAF344. The 3’homology arm was amplified with OAF345 and OAF346. The AID sequence was cloned from pLZ29 with OAF334 and OAF335. The backbone of plasmid containing the Self-Excising Selection Cassette was amplified in two fragments with OAF339/ OAF340 and OAF343/ OAF337 from pDD282.

To generate the *gei-17(zh142)* allele, an oligonucleotide corresponding to a target sequence near the *gei-17* translational start site (sgRNA: GTC GTT TCG AGA CAC AGC GG) was cloned into the plasmid pMW46. The 5’homology arm was amplified from genomic DNA with OAF336 and OAF338. The 3’homology arm was amplified with OAF341/ OAF342. The backbone containing the Self-Excising Selection Cassette and AID sequence was cloned in two fragments with OAF334/ OAF340 and OAF343 /OAF337 from pAF56, a previously cloned repair template for AID::SMO-1.

To generate the LIN-1 sumoylation site mutants *lin-1(zh157)* (K10A), *lin-1(158)* (K169A) and *lin-1(zh159*) (K10A, K169A), genome editing was performed according to the co-CRISPR strategy described by Arribere et al. (2014). To introduce the K10A mutation an oligonucleotide corresponding to a target sequence (sgRNA: GTC GAG TTC GGA AGA AGC CG) was cloned into plasmid pMW46. To introduce K169A mutation an oligonucleotide corresponding to a target sequence (sgRNA: GTT CAT ATT TGA GGA AAA GT) was cloned into the plasmid pMW46. The following constructs with indicated final concentration were microinjected into young adult *lin-1(st12212)* hermaphrodites: *dpy-10* sgRNA pJA58 (25 ng/μl), *dpy-10* repair oligonucleotide AF-ZF-827 (0.5 nM), *lin-1* sgRNA (75 ng/μl), *lin-1* repair oligonucleotide OAF377 (0.5 nM, introducing an NruI restriction site for K10A) or OAF378 (0.5 nM, introducing a SacII restriction site for K169A). To generate the *zh159* double mutant, the sgRNA#4 plasmid and OAF378 repair oligonucleotide was injected with the *dpy-10* sgRNA plasmid and *dpy-10* repair oligonucleotide into *lin-1(157)* hermaphrodites at the same concentrations as for the single mutant. Transformant showing a Rol phenotype were transferred to separate NGM plates, and animals containing the desired point mutations were identified by PCR amplification using the primers OAF365/ OAF366 for K10A or OAF367/ OAF368 for K169A, followed by restriction digests with NruI or SacII, respectively. The new *lin-1* alleles were sequenced and back-crossed three times to N2.

### Auxin treatment

NGM plates containing 1 mM auxin were prepared according to Zhang et al. (2015), seeded with OP50 *E. coli* bacteria and used immediately for the experiments. The auxin treatment protocol was adapted for each strain due to the differences in strain viablility and fertility. For strains containing AID-tagged GEI-17 (*gei-17(fgp1)* and *gei-17(zh142)* alleles), animals were synchronized by bleaching, and hatched L1 larvae were plated on auxin or control plates. Control plates contained the same dilution of ethanol, in which the auxin stock solution was prepared, as auxin plates. Animals were incubated at 25 °C and analyzed after 24 h or 36 h of treatment during the L3 or adult stage, respectively. Since homozygous *smo-1(zh140)* animals are sterile, they were maintained balanced with *tmC20*, and homozygous *smo-1(zh140)* animals were selected for the experiments. AID-tagged SMO-1, animals were likewise synchronized by bleaching, but hatched L1 larvae were first plated on standard NGM plates containing OP50 and incubated at 20 °C for 24 h, followed by transfer to auxin or control plates and 24 h of treatment. L3 animals were imaged right after treatment was complete, animals for analysis in the adult stage were instead transferred to standard NGM plates and analyzed 24 h later. For experiments involving different treatment periods, *smo-1(zh140)* animals were put on auxin/control plates 12 h, 24 h, 30 h and 36 h after L1, and transferred back to standard NGM plates 48 h after L1. Homozygous adults were analyzed 24 h later.

### Western blot analysis of the efficiency and kinetics of auxin-induced protein degradation

40 adult animals were transferred to an Eppendorf tube containing 20 μl of water. 20 μl of 2xSDS buffer were added and the sample was boiled for 5 min at 95 °C. In order to digest the DNA, 1 μl of DNase (Qiagen) was added, the sample was incubated for 5 min at room temperature and boiled again. Proteins were separated by SDS PAGE on 4-12% acrylamide gels and blotted onto PVDF membranes. After blocking non-specific binding sites with 5 % milk or bovine serum albumin in TBST (20 mM Tris, 150mM NaCl, 0.1 % Tween 20), the membranes were incubated with the primary antibody diluted in TBST containing 5 % milk overnight at 4 °C. After incubation with HRP-conjugated secondary antibodies, the protein bands were viualized by chemiluminescence using the SuperSignal West Pico or Dura Chemiluminescent Substrate (Thermo Scintific). Quantification was performed by measuring the band intensities using Fiji’s measurement tools (Schindelin et al. 2012). The following antibodies were used: anti-SUMO-1 1:500 (S5446 Sigma), anti-Flag 1:3000 (Sigma F3165-1MG), anti-Tubulin 1:10 000 (Abcam ab18251), HRPGoat anti-Rabbit 1:2000 (Jackson ImmunoReserach 111-035-144) and HRP Goat anti-Mouse 1:2000 (Jackson ImmunoReserach 115-035-146).

### Microscopy and image processing

For Nomarski and fluorescence imaging, live animals were mounted on 4% agarose pads and immobilized with 20 mM tetramisole hydrochloride solution in M9 buffer, unless stated otherwise. For toroid analysis in *lin-1* mutants, we used a custom microfluidic devices to immobilize the animals and performed imaging as described (Berger et al. 2021). Images were acquired with a Leica DM6000B microscope equipped with Nomarski and fluorescence optics, as well as a Leica DFC360FX camera and 63x (N.A. 1.32) oil immersion lens; a Leica DMRA microscope controlled by a custom build Matlab script, equipped with an image splitter and two Hamamatsu ORCA-flash 4.0L+ cameras to simultaneously acquire z-stacks in the DIC, mCherry and GFP channels using a 63x (N.A. 1.32) oil immersion lens; or a Matlab controlled Olympus BX61 microscope equipped with a X-light V2 spinning disc confocal system, a Prizmatix UHP-T-460-DI/UHP-T-560-DI LED as light source, an Andor iXon ultra888 EMCCD camera and a 60x (N.A 1.3) or 100x Plan Apo (N.A 1.4) oil immersion lens. Images were analyzed and quantified with Fiji software (Schindelin et al. 2012).

### Scoring vulval induction and morphogenesis

The numbers of induced VPCs was scored in synchronized L4 animals as described in Schmid et al. (2015). A score of 1 was assigned to a VPC when it underwent three division rounds and 0.5 when only one of the two VPC descendants had differentiated. A score of 0 was assigned to uninduced VPCs that had divided once and fused with the hypodermis.

Vulval lumen morphogenesis was assessed based on DIC microscopy at the L4 stage (L4.3-L4.7). Vulval defects in adult animals were scored by using a dissecting scope. Any abnormality in the vulval tissue visible under a dissecting microscope was categorized as ’abnormal vulva’ phenotype.

### Analysis of toroid formation

Animals at the L4 stage were imaged either on agar pads or in microfluidic devices (Berger et al. 2021) at 60x or 100x magnification, and z-stacks with a spacing of 0.13 to 0.2 μm were acquired. Toroid formation was monitored using the *swIs79[ajm-1::gfp]* or *cp21[hmr-1::gfp]* adherens junction markers (Diogon et al. 2007; Marston et al. 2016). Images were deconvolved either by the Huygens deconvolution software (Scientific Volume imaging) or using the Deconvolution lab plugin in Fiji (Schindelin et al. 2012). The measurement of the vulA and vulB1 diameters was done in xz-views of the cropped ventral toroids. The toroid fusion defects were scored in 3D reconstructed z-stacks.

### Quantification of LIN-1::GFP and EGL-17::YFP expression levels

Animals at the mid-L3 stage were imaged at 63x magnification using a wide-field microscope, acquiring z-stacks with a spacing of 0.3 μm. The average intensity of the nuclear LIN-1::GFP signal was measured in background subtracted, summed z-projections of 3 mid-sagittal sections of the VPCs. The nuclei of the 1° and 2° VPC descendants were manually selected, and the mean nuclear signal intensity was measured using the built-in measurement tools in Fiji. The data represent the averaged measurements for each VPC lineage (two nuclei at the Pn.px and four at the Pn.pxx stage). *egl-17::yfp* expression levels were analyzed in background subtracted mid-sagittal sections of the P6.x-P6.xxx cells. Cell bodies were manually selected, and the mean intensity was measured in Fiji.

### AC mispositioning and BM breaching shift analysis

Worms between L4.0-L4.5 were imaged at 63x or 100x magnification and z-stacks with a spacing of 0.1-0.3 μm were acquired. The AC position was monitored based on the *qyIs50[cdh-3>mCherry::moeABD; unc-119(+)]* reporter and DIC images, and the BM breach with the *qyIs10[lam-1>lam-1::gfp]* reporter. To assess the alignment of the AC with the 1° VPCs, the angle between a line through the middle of the vulval invagination and the center of the ACs nucleus and the dorso-ventral axis was measured, as illustrated in **Fig. 2D**. To quantify the BM breaching shift, the angles α and β between the middle of the vulval invagination to each of the BM breach points were measured and the ratios of the two angles was calculated (**Fig. 2D**).

### Statistical Analysis

Statistical analysis was performed using GraphPad Prism as indicated in the figure legends. Data were tested for parametric distribution and outliers were removed from analysis. For non-parametric continuous data, we used the Kolmogorov-Smirnov test, for non-continuous data (e.g. VPC induction counts) the Mann-Whitney test. Numerical values used for statistical analysis can be found in S1_Data excel file.

## Acknowledgements

We would like to thank all members of the Hajnal laboratory, Frauke Melchior, Damian Brunner and Ulrike Kutay for input, and the Caenorhabditis Genetics Center (funded by NIH Office of Research Infrastructure Programs (P40 OD010440)) for providing strains. This work was supported by grants from the Swiss National Science Foundation no. 31003A-166580 and the Swiss Cancer league no. 4377-02-2018 to AH.

## Author contributions

Aleksandra Fergin: Conceptualization, Data curation, Formal analysis, Investigation, Methodology, Visualization, Writing – original draft

Institute of Molecular Life Sciences, University of Zurich, Winterthurerstrasse, Zürich, Switzerland, Molecular Life Science PhD Program, University and ETH Zurich, Zürich

Gabriel Boesch: Investigation, Methodology

Institute of Molecular Life Sciences, University of Zurich, Winterthurerstrasse, Zürich, Switzerland, Molecular Life Science PhD Program, University and ETH Zurich, Zürich

Simon Berger: Writing – review & editing

Institute of Molecular Life Sciences, University of Zurich, Winterthurerstrasse, Zürich, Switzerland, Molecular Life Science PhD Program, University and ETH Zurich, Zürich

Nadja Greter: Investigation

Institute of Molecular Life Sciences, University of Zurich, Winterthurerstrasse, Zürich, Switzerland, Molecular Life Science PhD Program, University and ETH Zurich, Zürich

Alex Hajnal: Conceptualization, Investigation, Funding acquisition, Project administration, Supervision, Writing – review & editing

Institute of Molecular Life Sciences, University of Zurich, Winterthurerstrasse, Zürich, Switzerland

